# TOR signalling is required for host lipid metabolic remodelling and survival following enteric infection in *Drosophila*

**DOI:** 10.1101/2021.07.12.452110

**Authors:** Rujuta S. Deshpande, Byoungchun Lee, Yuemeng Qiao, Savraj S. Grewal

## Abstract

When infected by enteric pathogenic bacteria, animals need to initiate local and whole-body defence strategies. While most attention has focused on the role of innate immune anti-bacterial responses, less is known about how changes in host metabolism contribute to host defence. Using Drosophila as a model system, we identify induction of intestinal target-of-rapamycin (TOR) kinase signalling as a key adaptive metabolic response to enteric infection. We find TOR is induced independently of the IMD innate immune pathway, and functions together with IMD signalling to promote infection survival. These protective effects of TOR signalling are associated with re-modelling of host lipid metabolism. Thus, we see that TOR switches intestinal metabolism to lipolysis and fatty acid oxidation. In addition, TOR is required to limit excessive infection mediated wasting of adipose lipid stores by promoting an increase in the levels of fat body-expressed de novo lipid synthesis genes. Our data support a model in which induction of TOR represents a host tolerance response to counteract infection-mediated lipid wasting in order to promote survival.

## INTRODUCTION

Animals are constantly exposed to bacterial pathogens in their environment. As a result, they must be able to sense invading pathogens and then trigger appropriate defence responses. One defence strategy is to decrease pathogen load (Schneider and Ayres, 2008). Central to this mechanism are the innate immune responses. These are responsible for sensing invading bacteria at the sites of infection and then activating both local and whole-body host anti-bacterial responses (Buchon et al., 2014).

It also becoming clear that changes in host metabolism are another important defence strategy against infection (Ayres and Schneider, 2012; Medzhitov et al., 2012; Troha and Ayres, 2020). The innate immune response can be energetically costly, and these metabolic changes are often needed to fuel the immune response (Krejcova et al., 2019; Man et al., 2017). In addition, metabolic reprogramming is often essential for animals to adapt to and tolerate the presence of pathogens (Ganeshan et al., 2019; Sanchez et al., 2018; Wang et al., 2018; Wang et al., 2016; Weis et al., 2017). However, compared to our understanding of innate immunity, less is known about how metabolic adaptations promote host fitness upon infection.

Drosophila has provided a powerful model system to study host defence responses to enteric bacterial infection (Buchon et al., 2014; Lee and Lee, 2018). Upon ingestion of pathogenic bacteria, the intestine triggers two main responses to mount antibacterial defenses. The first involves activation of a the conserved IMD/Relish/NF-KappaB pathway by gram-negative bacteria, which leads to production of antimicrobial peptides (AMPs) (Buchon et al., 2014). The second involves bacteria-derived uracil, which stimulates reactive oxygen species (ROS) production in intestinal epithelial cells (Lee et al., 2018; Lee et al., 2015). Both pathways can promote local antimicrobial responses in the intestine and also trigger signalling from the intestine to other tissues to promote whole-body anti-bacterial response such as production of AMPs from the fat body (Wu et al., 2012; Yang et al., 2019). Enteric infection can also alter both intestinal and whole-body metabolism, but the contributions of these effects to defence against pathogens are not fully clear (Galenza and Foley, 2019; Lee and Lee, 2018; Wong et al., 2016).

TOR kinase is a conserved regulator of cell, tissue and whole-body metabolism (Ben-Sahra and Manning, 2017; Howell et al., 2013; Saxton and Sabatini, 2017). In general, TOR is activated under favourable conditions (e.g. growth factor stimulation and nutrient availability) to stimulate cellular anabolic metabolism and promote growth. In contrast, under stress conditions such as starvation, hypoxia or oxidative damage, TOR is inhibited to promote catabolic metabolism to ensure cell survival. The utility of *Drosophila* genetics has also been instrumental in showing how TOR activation in specific tissues can trigger and coordinate whole body-level physiological and metabolic responses (Boulan et al., 2015; Grewal, 2009; Texada et al., 2020). These effects rely on the ability of TOR signalling to promote inter-organ communication and endocrine signalling and have been shown to be essential for organismal responses to environmental changes such as altered nutrition and hypoxia (Boulan et al., 2015; Koyama et al., 2020).

Given the central role for TOR in controlling whole-body physiology and metabolism, some studies have begun to explore its role in responses to bacterial infection in *Drosophila*. However, these studies have differed in their conclusions about whether TOR activity is helpful or harmful to host immune responses and fitness upon infection. In some cases, it was reported that reduced TOR activity provided a benefit to the host. For example, enteric infection with *Ecc15*, a gram-negative bacterium, was shown to inhibit TOR and lead to increased lipid breakdown in the gut (Lee et al., 2018). This loss-of-TOR mediated metabolic shift to lipid catabolism was required for the antimicrobial ROS response and increased the host resistance to enteric infection. Similarly, another report showed that lowered TOR activity upon enteric infection could induce AMPs (Varma et al., 2014). Finally, one report showed that lowering TOR activity, either genetically or by nutrient restriction, was sufficient to increase survival upon systemic infection with either *P. aeruginosa* or *S. aureus* (Lee et al., 2017). In contrast to these findings, other studies showed that lowered TOR activity is detrimental to hosts upon infection. For example, enteric infection with *P*.*entomophila* decreased gut TOR activity, leading to suppressed intestinal protein synthesis, which reduced immune responses and prevented proper intestinal tissue repair (Chakrabarti et al., 2012). Another study also showed that TOR inhibition reduced fly survival upon systemic infection with *B. Cepacia* (Allen et al., 2016). The reasons for these difference in the links between TOR and infection response in Drosophila may be due to the different bacterial infections used or because of differences in host metabolic or nutrient status. Nevertheless, they indicate that further work is required to clarify how TOR may play a role in immune and metabolic responses to infection. We address this issue in this paper. We show that enteric infection leads to increased TOR signalling independently of innate signalling, and that this induction is required to remodel host lipid metabolism and promote survival.

## RESULTS

### Enteric bacterial infection stimulates TOR signalling in the adult intestine

TOR kinase couples environmental signals to changes in cellular metabolism. Generally, TOR has been shown to be activated by favorable conditions (e.g., abundance of nutrients and growth factors), while being inhibited by stress conditions (e.g., starvation, low oxygen, oxidative stress). We were interested in examining how TOR activity might be affected by enteric bacterial infection. We first infected flies with the gram-negative bacteria *Pseudomonas entomophila (P*.*e)* for 4hr and then dissected intestines for western blotting. Ribosomal protein S6 kinase (S6K), is directly phosphorylated and activated by TOR, hence we used western blotting for phosphorylated S6K as a readout for TOR activity. We found that oral *P*.*e*. feeding lead to increased phosphorylated S6K levels (Figure 1A). This increase was blocked by pre-feeding the flies rapamycin, a TOR inhibitor, indicating that the induction of phosphorylated S6K was through an increase in TOR activity (Figure 1B). We also examined phosphorylation of ribosomal protein S6, a downstream target of ribosomal protein S6 kinase. We saw that 4hrs of oral *P*.*e* infection also induced phosphorylated S6 levels in the intestine (Figure 1C). Moreover, when we performed immunostaining with the anti-phosphorylated S6 antibody, we saw that the increase in TOR activity was apparent in all cell types in the intestine, especially the large polyploid epithelial enterocytes that make up the bulk of the intestine (Figure 1D). A conserved function of TOR is the stimulation of cellular protein synthetic capacity, in large part mediated via upregulation of tRNA and rRNAs, and genes involved in ribosome synthesis (Ghosh et al., 2014; Killip and Grewal, 2012; Marshall et al., 2012; Mayer and Grummt, 2006; Rideout et al., 2012). When we used qRT-PCR to measure RNA levels in intestinal samples, we saw that oral *P*.*e*. infection led to an increase in tRNA and pre-rRNA levels and an increase in mRNA levels of three ribosome biogenesis genes, *Nop5, ppan, fibrillarin* upon oral *P*.*e* infection (Figure 1E). We explored these effects of oral bacterial infection further by performing a time course following oral *P*.*e*. feeding. We saw that the induction of TOR was rapid (within 4hrs of infection) and persisted for 24hrs of the oral infection period (Figure S1A). We also found that this induction of TOR was similar in males and females (Figure S1B). Moreover, the effects of *P*.*e*. appear limited to adults since 4hr oral infection in larvae didn’t increase phosphorylated S6K levels, and in fact showed a small decrease (Figure S1C). We also tested two other pathogenic gram-negative bacteria, *Vibrio cholera (V*.*c*.) and *Erwinia carotovora carotovora (Ecc15*). We again used western blotting for phosphorylated S6K to measure TOR and saw that oral infection with *V*.*c*. and *Ecc15* both led to increased intestinal TOR activity (Figure 2A). Together, these data indicate that induction of intestinal TOR kinase signalling is a rapid response to enteric gram-negative bacterial infection and that it stimulates the protein biosynthetic capacity of intestinal epithelial cells.

**Figure 1.**
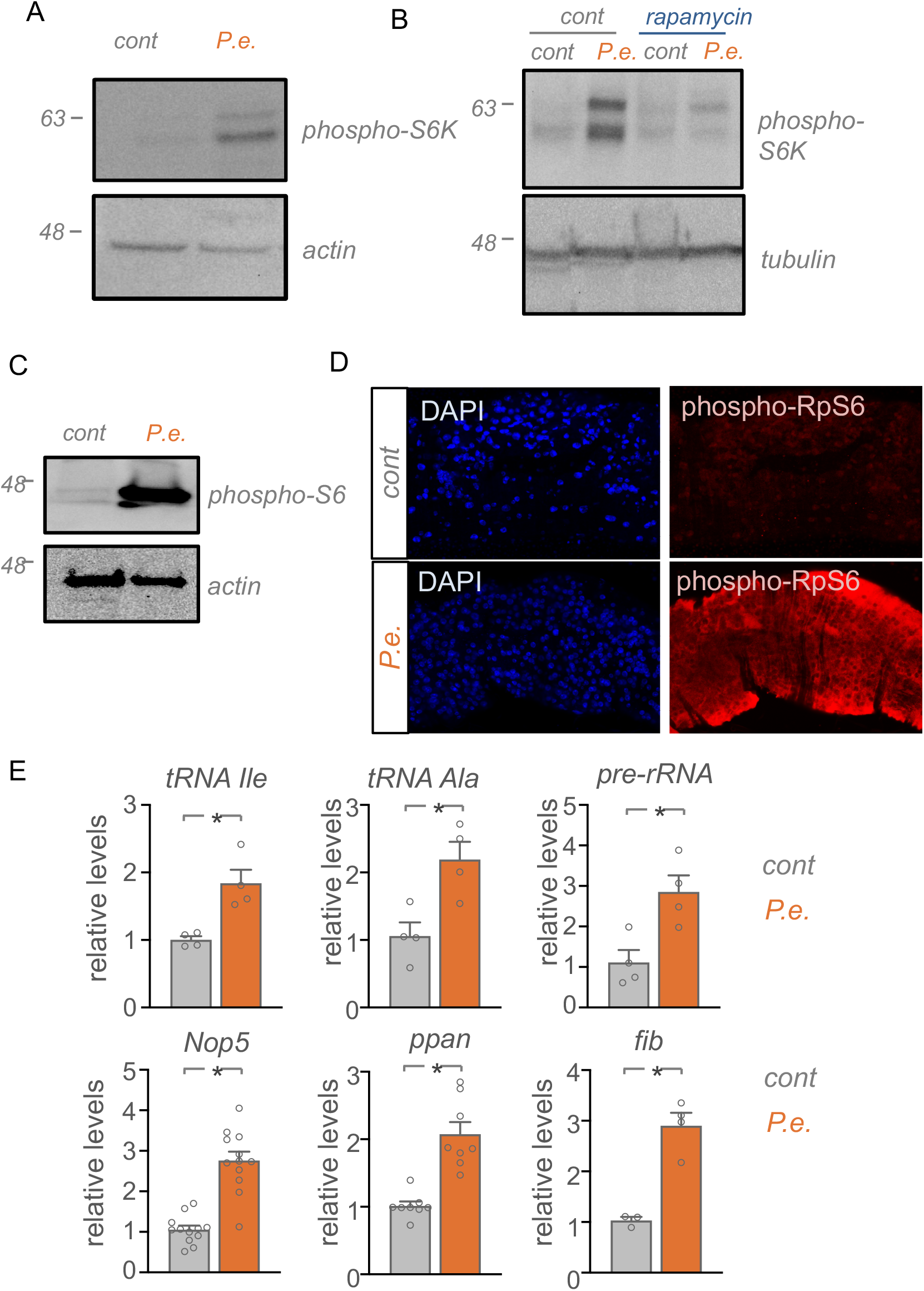
Enteric bacterial infection stimulates TOR activity in the intestine. A, B) Adult *w*^*1118*^ mated females were subjected to 4hr oral *P*.*e*. infection (B) or 24hr rapamycin pre-treatment followed by 4hr oral *P*.*e*. feeding (B). Dissected intestines were lysed and analyzed by western blotting using antibodies to phosphorylated*-*S6K and actin or tubulin (shown as loading controls). C, D) Adult *w*^*1118*^ mated females were subjected to 4hr oral *P*.*e*. infection. Intestines were then either lysed and processed for western blotting using antibodies to phosphorylated ribosomal protein S6 and actin, (C) or stained with phosphorylated-S6 antibody for immunofluorescence, (D). Blue= Hoechst DNA dye; red= phosphorylated-S6 antibody. E) Adult *w*^*1118*^ mated females were subjected to 4hr oral *P*.*e*. infection. Dissected intestines were processed for qRT-PCR analysis for the indicated RNAs. Data are represented as bar graphs with error bars indicating the S.E.M. *p<0.05, Student’s Unpaired t-test.

**Figure 2.**
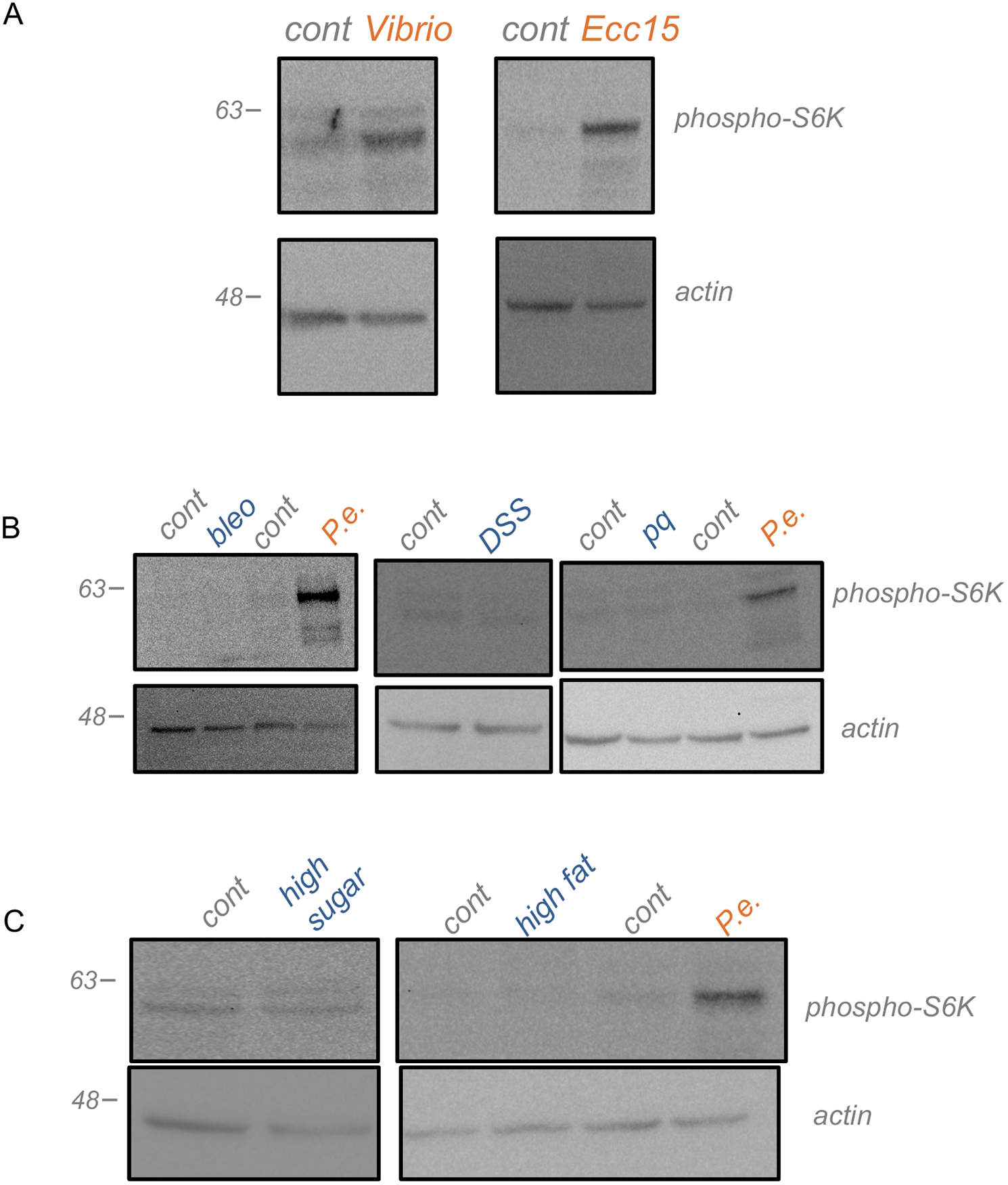
Enteric bacterial infection, but not other environmental stress, stimulates TOR activity in the intestine. **A)** Adult *w*^*1118*^ mated females were subjected to 4hr oral infection treatments of pathogenic gram-negative bacteria, *Vibrio cholera (V*.*c*.) and *Erwinia carotovora carotovora (Ecc15*). Dissected intestines were lysed and processed for western blotting using antibodies against phosphorylated-S6K and actin (loading control). B) Adult *w*^*1118*^ mated females subjected to 4hr treatments of chemical stressors: 25μg/ml Bleomycin, 5%DSS, 2mM paraquat. Dissected intestines were lysed and processed for western blotting using antibodies against phosphorylated-S6K and actin (loading control). A representative blot is shown. C) Adult *w*^*1118*^ mated females subjected to 4hr treatments of nutrient stresses high sugar (40% sucrose) (left) and high fat (30% lard) (middle) diets, or Pe infection (positive control). Dissected intestines were lysed and processed for western blotting using antibodies against phosphorylated-S6K and actin (loading control).

Studies in *Drosophila* and mammalian cells have shown that two conserved signalling pathways-PI3K/AKT and extracellular-regulated kinase (ERK) pathway can promote TOR signalling (Shaw and Cantley, 2006). Moreover, both pathways have been shown to mediate effects on intestines such as growth and proliferation in response to stimuli such as infection and starvation downstream of growth factor signalling (insulins and EGFs)(Miguel-Aliaga et al., 2018). We therefore performed western blotting on infected and control intestines for phosphorylated AKT (PI3K/AKT pathway) and phosphorylated ERK (ERK pathway). We found that oral *P*.*e*. had no effect on PI3K/AKT or ERK pathway, suggesting that *P*.*e*. effects may be specifically inducing TOR activation (Figure S1D). We next explored whether other enteric intestine stresses might also regulate intestine TOR signalling. Feeding flies with three known chemical intestine stressors, bleomycin (a DNA damaging agent), dextran sodium sulphate (a detergent) and paraquat (an oxidative stressor), had no effect on intestine TOR activity (Figure 2B). We also explored two nutrient stresses - high sugar and high fat. However, we saw that feeding the flies either a high sugar (40%) or high fat (30%) supplemented diet, also did not have any effect on TOR signalling in the intestine (Figure 2C). Together our data suggests that the induction of TOR appears specific to oral bacterial infection.

### Infection mediated TOR stimulation is independent of IMD and ROS signalling

We next investigated how infection might stimulate TOR. Two primary and well-studied responses to enteric gram-negative bacterial infection are activation of the Immune deficiency (IMD)/ NF-kB pathway (Kleino and Silverman, 2014) and the induction of reactive oxygen species (ROS) in the intestinal enterocytes (Lee et al., 2015; Wu et al., 2012). We therefore examined whether either IMD or ROS activation trigger TOR induction upon oral *P*.*e*. infection. The IMD signalling cascade stimulates the NF-ºB-like transcription factor, Relish, to induce an anti-bacterial immune response to gram-negative infections. We began by testing the involvement of IMD signalling by using mutants for Imd and Relish, two components of the pathway. We infected either control (*w*^*1118*^) or either *rel* or *imd* mutants with *P*.*e*. for 4hr and then measured intestinal phosphor S6K levels. We found that the induction of intestinal TOR upon oral *P*.*e* infection was still observed in both the *relish* and *imd* mutants suggesting that TOR induction is independent of IMD signalling (Figure 3A, B).

**Figure 3.**
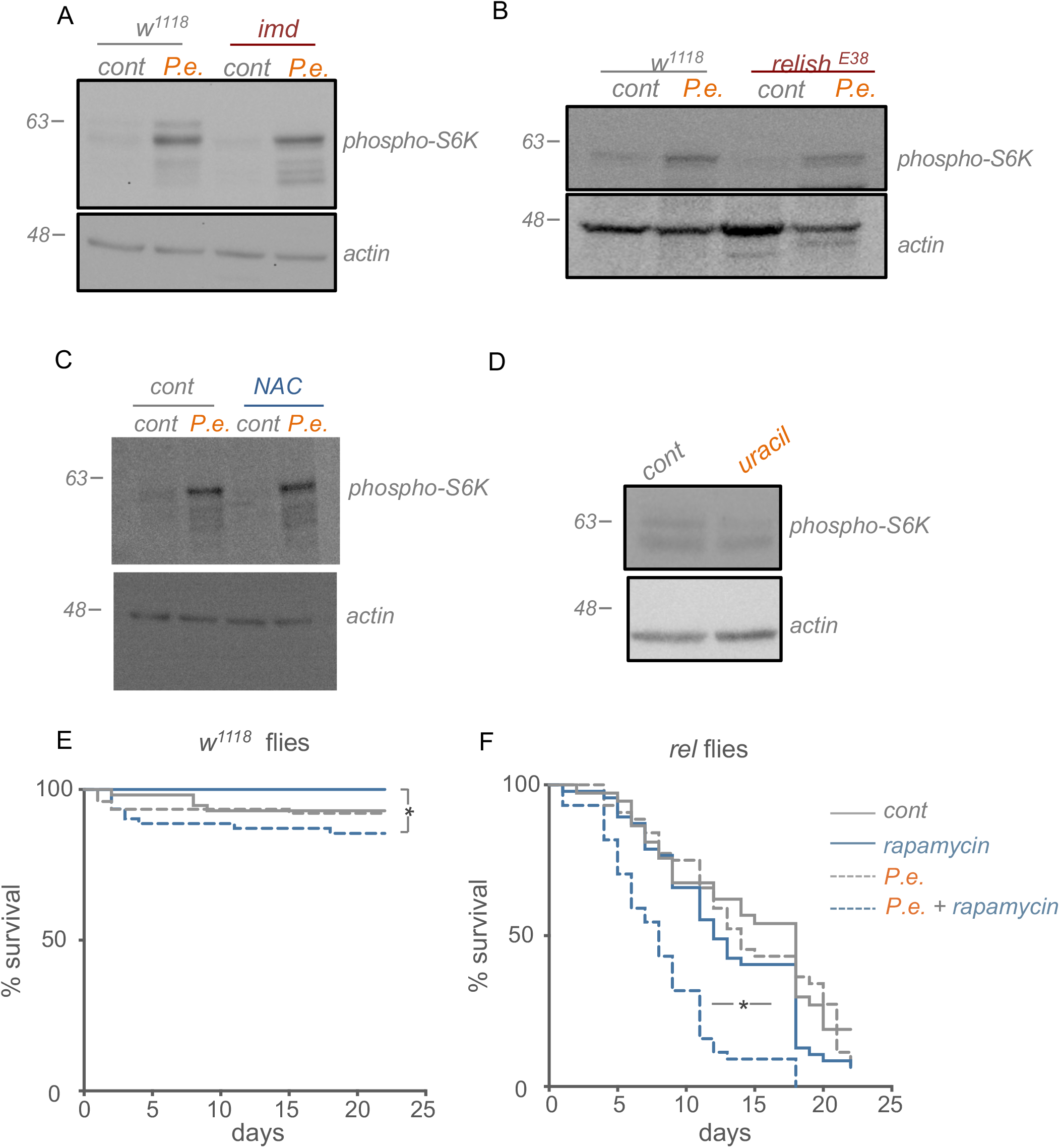
TOR and IMD signaling function in parallel to control survival in response to enteric infection. A) Adult *w*^*1118*^ and Immune deficiency (*imd*) mutants subjected to 4hrs oral *P*.*e*. infection. Dissected intestines were lysed and processed for western blotting using antibodies against phosphorylated-S6K and *actin* (loading control). **b** Adult *w*^*1118*^ and Rel/NF-ºB transcription factor *relish* mutants subjected to 4hrs oral *P*.*e*. infection. Dissected intestines were lysed and processed for western blotting using antibodies against phosphorylated-S6K and *actin* (loading control). C) Adult *w*^*1118*^ mated females subjected to feeding for 4hrs with *P*.*e*. alone (left) and *P*.*e*. + antioxidant N-acetyl cysteine, NAC (right). Dissected intestines were lysed and processed for western blotting using antibodies against phosphorylated-S6K levels and *actin* (loading control). D) Adult *w*^*1118*^ mated females subjected to 4hr uracil feeding. Dissected intestines were lysed and processed for western blotting using antibodies against phosphorylated-S6K levels and actin (loading control). E) Survival plot of control *w*^*1118*^ (E) and *relish*^*E20*^ (*rel*) mutant (F) mated female flies subjected to 48-hr oral *P*.*e*. infection. Animals were then returned to standard food and the percentage of animals surviving was counted. N = at least 50 animals per experimental condition. *p< 0.05, log rank test.

Next, we determined if ROS signalling plays a role in TOR induction upon infection. To do this we first examined the effects of blocking ROS using the antioxidant N-acetylcysteine (NAC). However, we found that feeding the flies a potent antioxidant, along with oral *P*.*e*. feeding didn’t reverse the induction of TOR upon infection (Figure 3C). Activation of NADPH dual oxidase (DUOX) is one of the immediate signalling events that triggers ROS activation and subsequent downstream anti-microbial responses upon enteric infection (Lee et al., 2015). Uracil is a bacterial derived compound that activates DUOX production and has been shown to induce both local and systemic responses to promote host survival upon infection (Lee et al., 2018). To determine whether TOR induction is downstream of the ROS pathway, we first subjected adult female flies to 4hrs of uracil feeding and found that uracil did not induce TOR signalling in the intestine (Figure 3D). Together with our result with the paraquat, a stimulator of ROS, our data suggests that TOR induction is independent of ROS, and suggest that it is activated in parallel to the well-described IMD/Relish and ROS pathways

### Inhibiting TOR and IMD pathways simultaneously reduces survival upon *P*.*e*. infection

We next examined the consequences of TOR induction upon *P*.*e*. infection. We first examined effects on survival following enteric *P*.*e*. infection. Under our laboratory fly culture conditions, the strain of *P*.*e*. we use is not strongly pathogenic. Thus, when we infected control (*w*^*1118*^) adult flies for 2 days and then monitored their survival over approximately three weeks, we saw little effect on viability compared to uninfected flies (Figure 3E). When we infected flies and simultaneously inhibited TOR by feeding flies rapamycin, we found that this induced a slight, but significant, decrease in survival compared to flies fed rapamycin alone (Figure 3E). We next tested the possibility that TOR functions in parallel IMD/Relish signalling to promote infection survival. We found that *relish* mutants had a generally reduced lifespan on our normal lab food compared to control (*w*^*1118*^) adults (Figure 3F). Either enteric infection with *P*.*e*., or blocking TOR with rapamycin, had no effect on viability in the *relish* mutants alone. However, when we infected *relish* mutants and simultaneously fed them rapamycin to inhibit TOR, we saw a significant decrease in survival compared to *relish* mutant flies subjected to infection or rapamycin treatment alone (Figure 3F). These results suggest that in order to survive enteric infection, an animal needs cooperative activation of both IMD/Relish and TOR signalling.

A primary infection response induced by the IMD pathway in flies is the production of anti-microbial peptides (AMPs) (Kleino and Silverman, 2014). We saw that oral infection with *P*.*e*. led to a strong induction of several AMPs, including *Cecropin A (CecA), Cecropin C (CecC), Metchnikowin (Mtk)*, and *Drosocin (Dro)* (Figure 4). However, when we inhibited TOR by feeding flies rapamycin this induction was not significantly affected (Figure 4). This suggests that the induction of TOR upon infection may not be required to induce antibacterial resistance responses, and that the requirement for TOR in infection survival may reflect a role in other immune responses.

**Figure 4.**
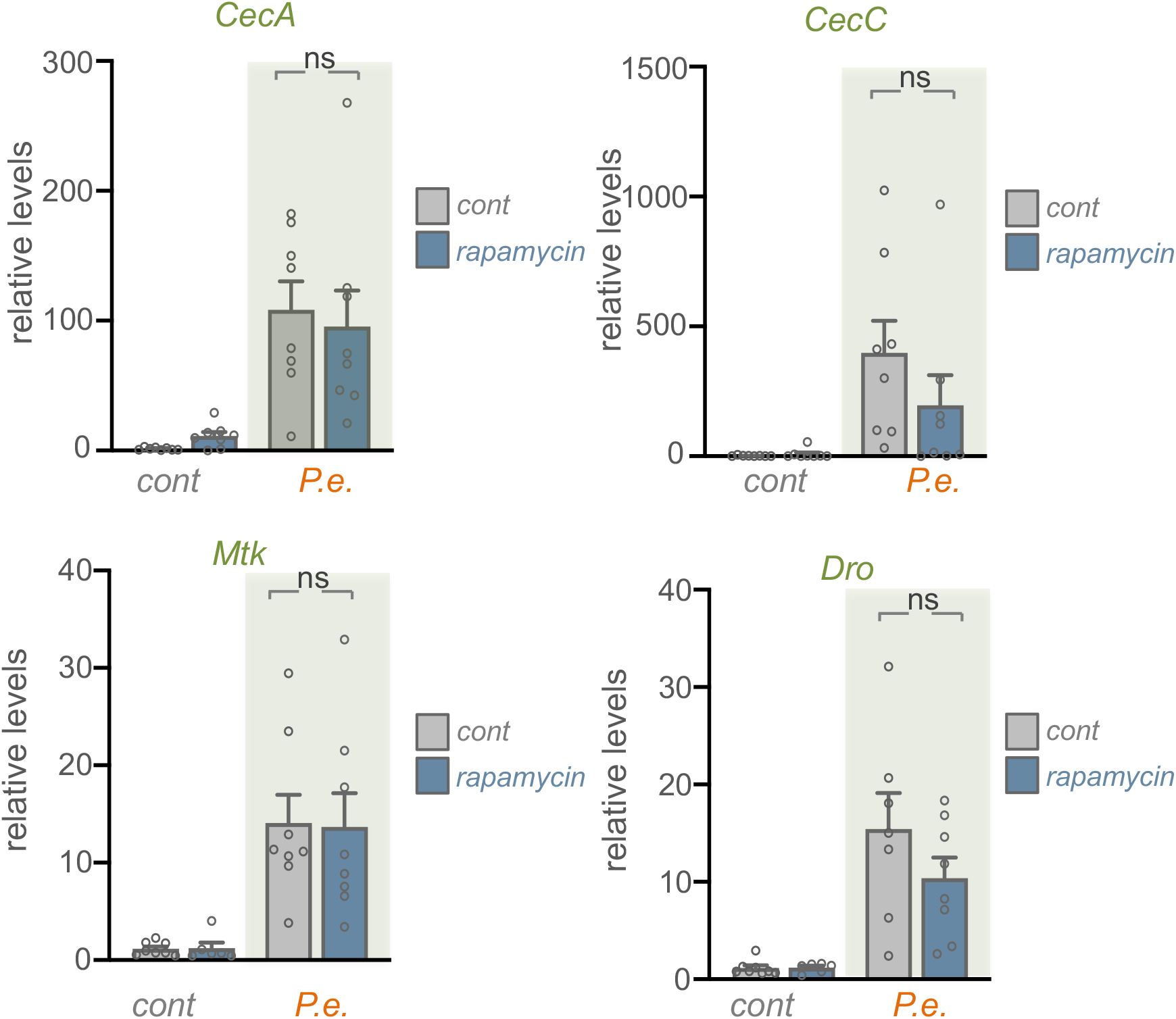
Induction of intestinal TOR signaling is not required for systemic AMP induction. qRT-PCR analysis on adult *w*^*1118*^ mated females subjected to a 24hr pre-treatment of rapamycin or DMSO control followed by 24hr oral *P*.*e*. feeding along with rapamycin. mRNA transcript levels of anti-microbial peptides (AMPs) are presented as relative changes vs control (corrected for RpS9). The bars represent the mean for each condition, with error bars representing the S.E.M and individual values plotted as symbols. ns = not significant, two-way ANOVA followed by Students t-test,

### TOR induction limits lipid depletion and promote lipid synthesis upon infection

Initiating immune responses against infection can be an energetically costly for infected hosts. Remodelling of host metabolism is therefore increasingly recognized as an important component of immune responses (Troha and Ayres, 2020). Given that TOR kinase is a conserved regulator of metabolism, we investigated whether it might play a role in modulating host metabolic responses to infection. Lipid stores are an important metabolic fuel source. Lipids can be synthesized and stored as triacyglycerides (TAGs) in lipid droplets in the fly fat body and oenocytes. They can be then mobilized, transported to tissues, and used to fuel metabolism, particularly in stress conditions (Heier and Kuhnlein, 2018). We tested for changes in TAGs upon oral *P*.*e*. infection and found a significant decrease in infected *w*^*1118*^ adult females compared to control flies (Figure 5A) as has been reported following systemic infection in flies (Chambers et al., 2012; Dionne et al., 2006). Since the majority of the lipid stores are stored in the fat body, we dissected the fat bodies of infected females, and found a decrease in the lipid droplet size by BODIPY staining (Figure 5B). In addition, we saw that enteric infection increased whole-body expression of two lipid binding proteins, apoLpp and Mtp, that are highly expressed in the fat body and that are needed for transport of lipids through the hemolymph (Figure 5C). These results suggest that infection leads to mobilization and transport of fat body lipid stores, perhaps as a way to provide lipids to other tissues to fuel their metabolism

**Figure 5.**
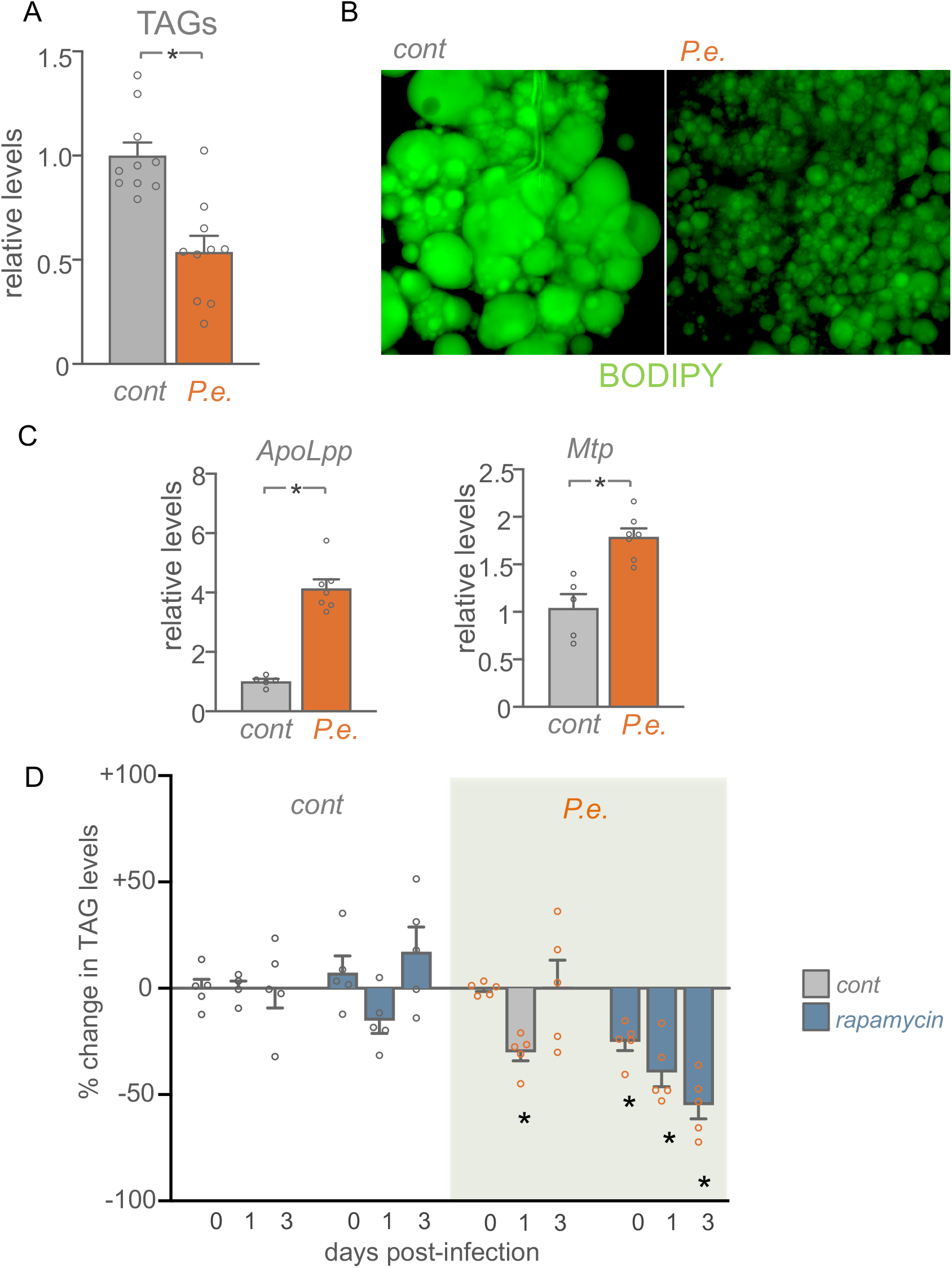
A) Adult *w*^*1118*^ mated females subjected to 24hr oral *P*.*e*. infection. Female flies were snap frozen on dry ice and TAG assays were performed. The bars represent percentage change in TAG levels (compared to uninfected control animals), normalized to the protein content for each condition. The bars represent the mean for each condition, with error bars representing the S.E.M. and individual values plotted as symbols. *p<0.05, Students t-test. B) BODIPY staining of fat body of *w*^*1118*^ *w*^*1118*^control and 24hr *P*.*e*. infected mated females. Green=BODIPY. C) Adult *w*^*1118*^ mated females subjected to 24hr oral *P*.*e*. infection. Whole animals were processed for qRT-PCR analysis for the indicated mRNAs. Data are represented as bar graphs with error bars indicating the S.E.M. p<0.05, Student’s Unpaired t-test. D) Adult *w*^*1118*^ mated females subjected to 24hr pre-treatment of rapamycin followed by 24hr oral *P*.*e*. feeding along with rapamycin. TAG assays were performed on flies at 0, 1 or 3 days after infection. The bars represent percentage change in TAG levels (compared to uninfected control animals), normalized to the protein content for each condition. The bars represent the mean for each condition, with error bars representing the S.E.M. and individual values plotted as symbols. * represents P <0.05 for each experimental group compared to the control group at the same timepoint.

We next examined what role TOR might play in these lipid effects by measuring TAG levels at 0-, 1- and 3-days following infection in control vs rapamycin-treated flies. In control flies, infection led to a transient decrease in TAG levels at the 1-day timepoint, but then TAGs recovered to the same level as uninfected flies at 3 days (Figure 5D). Rapamycin treatment alone had no significant effect on TAG levels at any timepoint compared to uninfected control flies (Figure 5D). However, when we infected flies and simultaneously fed them rapamycin to inhibit TOR, we saw a progressive depletion of TAG stores at each timepoint following infection (Figure 5D). These results suggest TOR is needed to limit excessive loss of lipid stores following infection. To do this, TOR may be blocking excess lipase function (to limit lipolysis) or may be increasing lipid synthesis (to resupply new lipids). We found that infected flies showed a significant upregulation in mRNA expression levels of genes required for de-novo lipid synthesis such as acetyl-CoA carboxylase (ACC), fatty acid synthetase 1 (FASN1), midway (mdy/DGAT1), dgat2, lipin, and lsd2 (Figure 6). Moreover, the expression levels of two transcription factors, SREBP and Mondo, which promote the transcription of these genes (Heier and Kuhnlein, 2018; Mattila and Hietakangas, 2017) were also upregulated (Figure 6). Interestingly, rapamycin treatment blocked the *P*.*e*.-induced increase in expression of these lipid synthesis genes and transcription factors (Figure 6). These results suggest that one role for the increased TOR activity that we see upon infection may be to induce de novo lipid synthesis to counteract the infection-mediated depletion of lipid stores.

**Figure 6.**
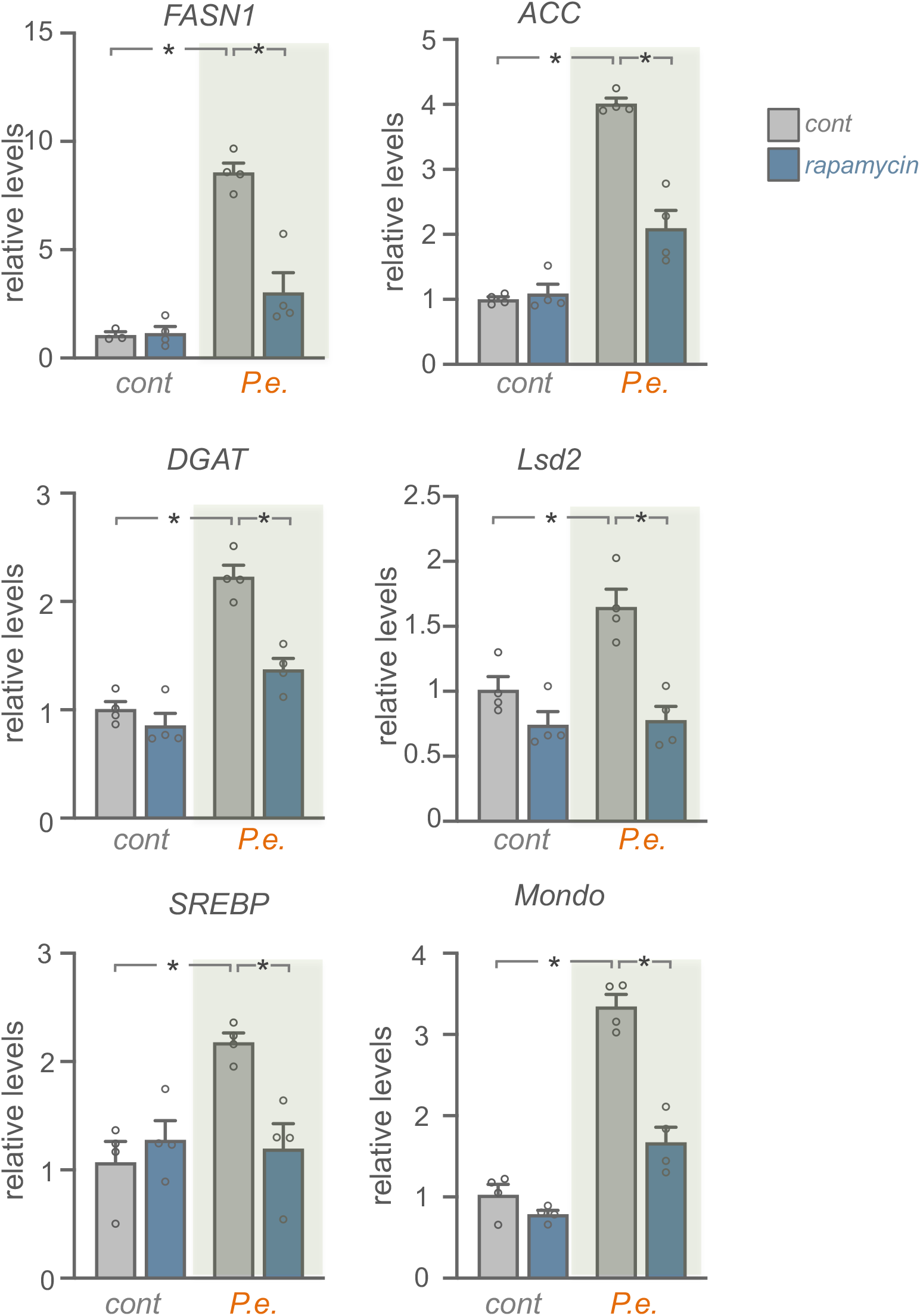
TOR is required for whole-body lipid synthesis gene induction upon enteric infection. A. qRT-PCR analysis of lipid synthesis genes and the transcription factors, SREBP and Mondo, in *w*^*1118*^ mated females pretreated for 24n hour with either DMSO (control) or rapamycin, followed by 24hr of either sucrose (control) or 24hr oral *P*.*e*. feeding (grey bars). The bars represent the mean for each condition, with error bars representing the S.E.M and individual values plotted as symbols. * p<0.05, two-way ANOVA followed by Students t-test.

### Infection promotes glycogen mobilization through TOR signalling

De novo lipid synthesis often relies on metabolic conversion of glucose into acetyl-CoA which can then serve as the source for new TAG synthesis (Heier and Kuhnlein, 2018). Flies can acquire glucose both from their diet and also from mobilization of stored glucose in the form of glycogen (Mattila and Hietakangas, 2017). When we infected flies with *P*.*e*. we saw a significant decrease in whole-body glycogen levels, as has been reported following systemic infection (Chambers et al., 2012; Dionne et al., 2006), and an increase in mRNA expression levels of several genes required for glycogen mobilization such as glycogen phosphorylase (GlyP), AGL/ CG9485, and UDP-glucose pyrophosphorylase (UGP) (Figure 7A, B). These results indicate that infection triggers a mobilization of host glycogen stores. Interestingly, when we blocked TOR signalling by feeding flies rapamycin, this prevented the infection induced increase in mRNA levels of GlyP, the limiting enzyme for glycogen mobilization (Figure 7C). Moreover, we saw that infection-mediated mobilization of glycogen was reduced in rapamycin-fed animals (Figure 7D). Taken together, these results suggest that infection leads to depletion of stored glycogen in part through TOR signalling. Thus, one possibility is that this TOR-dependent mobilization of glycogen is used to provide glucose for the TOR-induced de novo synthesis of TAGs.

**Figure 7.**
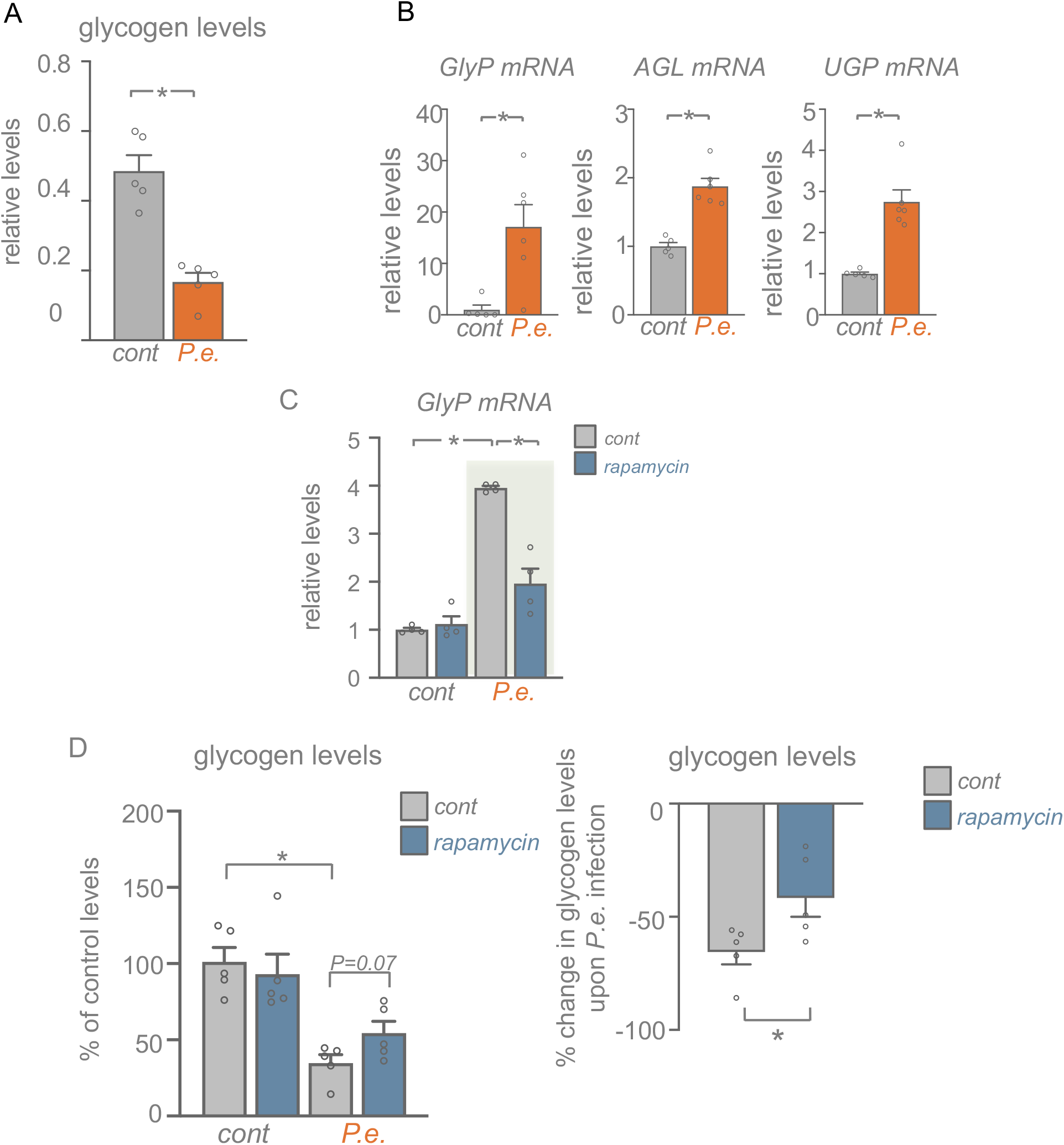
Enteric infection leads to glycogen mobilization in part through TOR activity. A) Adult *w*^*1118*^ mated females were subjected to 24hr oral *P*.*e*. infection and then whole-body glycogen levels were measured. The bars represent the mean for each condition, with error bars representing the S.E.M. and individual values plotted as symbols. * p<0.05, Students t-test. B) *w*^*1118*^ mated females subjected to 24hr of either sucrose (control) or 24hr oral *P*.*e*. feeding (orange bars) and then processed for qRT-PCR analysis of genes involved in glycogen breakdown. The bars represent the mean for each condition, with error bars representing the S.E.M. and individual values plotted as symbols. * p<0.05, Students t-test. C) *w*^*1118*^ mated females were pretreated for 24 hours with either DMSO (control) or rapamycin, followed by 24hr of either sucrose (control) or 24hr oral *P*.*e*. feeding (grey bars). Whole animals were then processed for qRT-PCR analysis of GlyP mRNA. The bars represent the mean for each condition, with error bars representing the S.E.M and individual values plotted as symbols. *, p<0.05, two-way ANOVA followed by Students t-test. C) *w*^*1118*^ mated females were pretreated for 24 hours with either DMSO (control) or rapamycin, followed by 24hr of either sucrose or 24hr oral *P*.*e*. feeding. Whole animals were then processed for measurement of total glycogen assays. Left, the bars represent the mean for each condition, with error bars representing the S.E.M. and individual values plotted as symbols. * p<0.05, two-way ANOVA, followed by Students t-test. Right, the data are presented as the percentage decrease in whole-body glycogen levels upon infection in control vs. rapamycin-treated samples. The bars represent the mean for each condition, with error bars representing the S.E.M. and individual values plotted as symbols. *p<0.05, Students t-test.

### Infection promotes TOR dependent intestinal lipid metabolism remodelling

Previous studies have shown that local changes in intestinal lipids can impact whole body lipid metabolism (Zhao and Karpac, 2020). Given that we identified TOR stimulation in the intestine following infection, we examined whether this might alter gut lipids. As has been reported previously (Kamareddine et al., 2018; Luis et al., 2016; Miguel-Aliaga et al., 2018), we saw that enterocytes in the anterior region of the midgut accumulated lipid droplets as visualized by Oil Red O and BODIPY staining. We found that oral infection with *P*.*e*. lead to a depletion of these intestinal lipids(Figure 8A, B). We also saw increased intestinal expression of three lipases, *brummer (bmm), CG5966*, and *Lipase 3 (Lip3)*, suggesting that infection depletes intestinal lipids through increase lipolysis (Figure 8C). Interestingly, this infection-mediated increased in gut lipase expression was prevented by rapamycin feeding suggesting that they were induced by the increased intestinal TOR signalling induced by enteric *P*.*e*. (Figure 8C). Mobilized lipids are often used to fuel beta-oxidation as an alternate to using glucose to fuel mitochondrial metabolism. This type of metabolism is often seen in host cells and tissues following pathogenic infection. When we infected flies with *P*.*e*. we saw increased expression of several genes involved in fatty acid beta-oxidation (*carnitine palmitoyltransferase 2 (CPT2), Carnitine O - Acetly Transferase (CRAT), Acy l-coenzyme A oxidase (Acox57Dp), Medium-chain acyl-CoA dehydrogenase (Mcad), ACSL1 (CG3961)* and *Acetyl Coenzyme A synthase (AcCoAS)*), and as with the lipases, these effects were prevented by rapamycin feeding (Figure 8D). These results therefore suggest that enteric pathogen infection leads to a TOR-dependent remodelling of intestinal metabolism toward lipid mobilization and beta oxidation.

**Figure 8.**
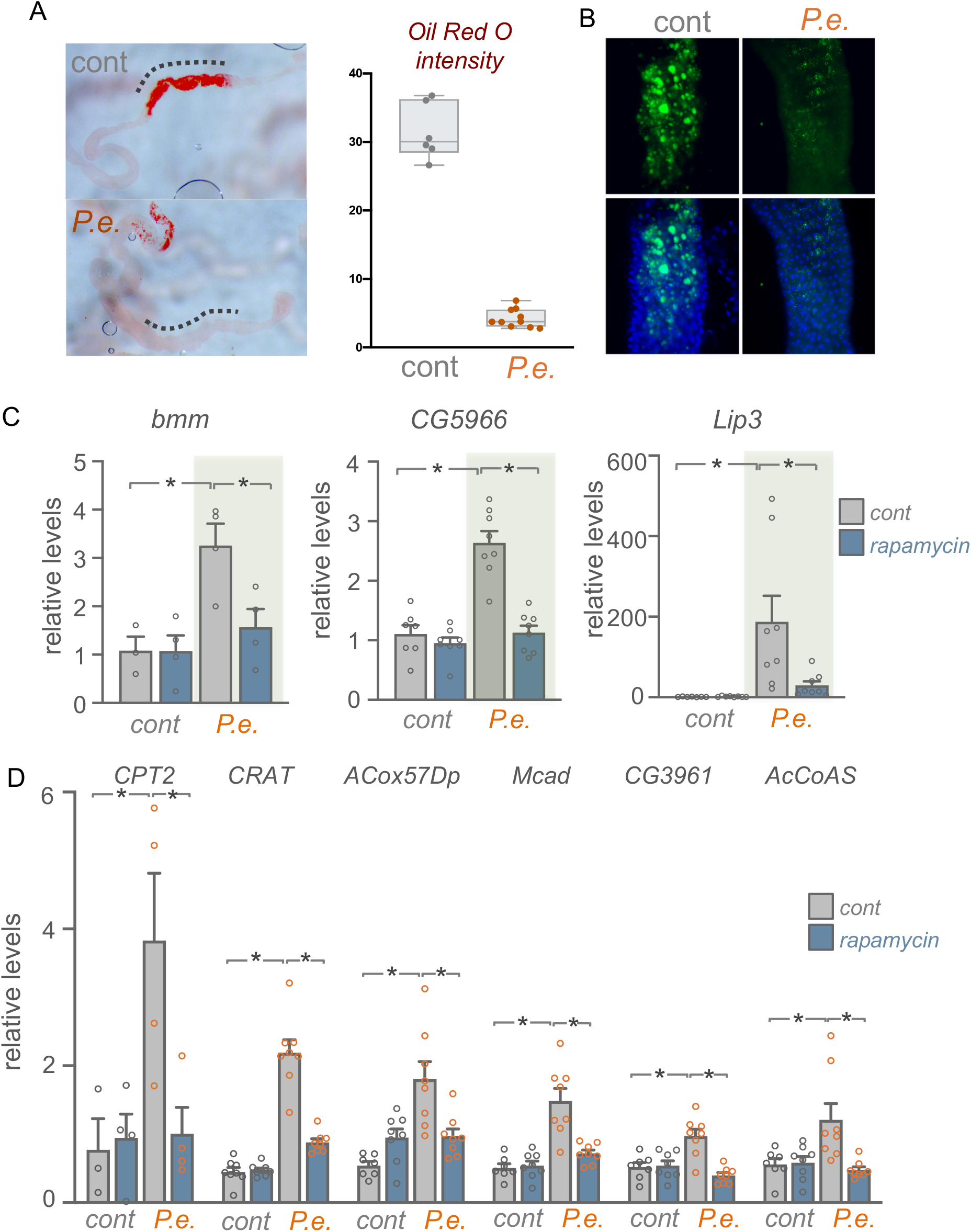
Infection promotes TOR-dependent intestinal lipid metabolism remodelling. A) Lipid droplet accumulation in the anterior region of the intestines stained with left, Oil-Red O (ORO) dye. The ORO intensities in the anterior regions (indicated with dash line) of *w*^*1118*^control and 24hr *P*.*e*. infected intestines were measured and presented as bar graphs representing mean for each condition, with error bars representing the S.E.M. and individual values plotted as symbols. * p<0.05, Students t-test. B) BODIPY staining of anterior regions of *w*^*1118*^control and 24hr *P*.*e*. infected intestines. Green=BODIPY, blue= Hoechst DNA dye. C, D) *w*^*1118*^ mated females were pretreated for 24 hours with either DMSO (control) or rapamycin, followed by 24hr of either sucrose (control) or 24hr oral *P*.*e*. feeding (grey bars). Whole animals were then processed for qRT-PCR analysis of lipase mRNAs (C) and mRNAs of genes involved in fatty acid beta-oxidation (The bars represent the mean for each condition, with error bars representing the S.E.M and individual values plotted as symbols. * p<0.05, two-way ANOVA followed by Students t-test.

## DISCUSSION

In this paper we show that enteric infection with gram-negative bacteria leads to increased intestinal TOR signalling, a response that functions together with induction of the IMD innate signalling pathway to promote infection survival. The induction of TOR occurred rapidly and was seen predominantly in the large enterocytes, the main absorptive, barrier and metabolic cells of the intestine. The mechanism by which infection stimulates TOR remains to be determined. We showed that it was independent of induction of ROS and IMD signalling, the two main pathways induced by gram-negative bacterial infection in the fly intestine. It is possible that a bacterial-derived secreted metabolite or small molecule may be responsible for stimulating TOR. This type of induction could also rely on pathogen interactions with commensal bacteria as such crosstalk has been shown to alter bacterial small molecule and metabolite secretion in the intestine and can affect host epithelial responses (McCarville et al., 2020).

We found that the TOR induction was not required for induction of the AMPs the main anti-bacterial resistance response in flies. The AMPs are primarily induced by IMD/Relish signalling following enteric infection. Interestingly we saw the infection survival was reduced only when we simultaneously blocked both IMD signalling (*relish* mutants) and TOR signalling (rapamycin feeding). Based on these data, one simple model is that upon infection, the IMD pathway is induced to initiate resistance (antibacterial defences), while TOR induction plays a role in tolerance responses (adaption to pathogen infection).

Tolerance responses are defined as alterations in host biology that limit pathology and promote survival without affecting pathogen load (Ayres and Schneider, 2012; Medzhitov et al., 2012). These can involve adaptations in host metabolism (Cumnock et al., 2018), changes in host behaviour, or induction of host tissue protective and repair processes (Martins et al., 2019). It is becoming clear that these responses are as important as resistance (anti-pathogenic) responses in determining host survival upon infection, and, as a result, there is increasing interest in determining mechanisms that control tolerance. In the context of enteric infection, recent studies have emphasized how gut-mediated changes in whole-body level physiological programs such as systemic insulin signalling and glucose and lipid metabolism play an important role in tolerance responses (Sanchez et al., 2018; Schieber et al., 2015). We saw that enteric infection led to a transient reduction in total TAGs and increased expression of lipoproteins. These results are consistent with an infection-mediated mobilization and transport of fat body-derived lipids to other tissues, perhaps to support their metabolism. Indeed, previous work has described how activation of the IMD pathway in the Drosophila fat body can promote lipid mobilization (Davoodi et al., 2019). Moreover, a switch to fatty acid oxidation is often a type of metabolic reprogramming seen upon infection (Cumnock et al., 2018).

In the context of infection-mediated lipid mobilization, we saw that the main function for TOR appeared to be limit excess lipid loss. Thus when we rapamycin-treated flies we saw that the transient decrease in lipid stores following infection developed into a progressive loss of lipid stores. Our results suggest that TOR functions to prevent excess lipid loss by promoting de novo lipid synthesis by increasing expression of lipid synthesis genes. These genes are enriched for expression in the fat body (Leader et al., 2018) indicating that enteric infection induces a TOR-dependent gut-to-fat body remote control of lipid synthesis. One way that this may happen is as a result of TOR-mediated change in intestinal lipid metabolism. We saw a TOR-dependent increase in lipolysis and beta oxidation, and previous work has shown that changes in intestinal lipid metabolism, particularly induction of lipolysis, can promote increases in whole-body TAG levels (Kamareddine et al., 2018; Karpac et al., 2013; Song et al., 2014; Zhao and Karpac, 2020). In addition, changes in intestinal lipid metabolism and lipolysis have been shown to alter other organismal phenotypes such as feeding and aging (Bouagnon et al., 2019; Luis et al., 2016).

A central theme of our work is that alterations in host lipid metabolism are important component of immune responses. This is supported by previous studies in flies that have described how both intestinal and fat body lipid metabolism are needed for effective immune responses (Chakrabarti et al., 2014; Harsh et al., 2019; Kamareddine et al., 2018; Lee et al., 2018; Martinez et al., 2020). We pinpoint TOR as a central modulator of enteric infection-mediated changes in lipid metabolism, likely as a mechanism of infection tolerance. The intestine also plays a central role coordinating other aspects of fly physiology such as repair of local tissue damage (Colombani and Andersen, 2020) and modulation of feeding behavior (Hadjieconomou et al., 2020; Miguel-Aliaga et al., 2018; Redhai et al., 2020). Given previous work implicating these processes as regulators of infection tolerance (Ayres and Schneider, 2012; Rao et al., 2017), our finding that TOR is induced in gut suggest it may also play a role in these other important responses to infection.

## METHODS

### Drosophila stocks and culturing

Flies were kept on medium containing 150 g agar, 1600 g cornmeal, 770 g Torula yeast, 675 g sucrose, 2340 g D-glucose, 240 ml acid mixture (propionic acid/phosphoric acid) per 34 L water and maintained at 25°C. The following lines were used in this study: *w*^*1118*^, *imd*^*[EY08573*^*], rel*^*E20*^, *rel*^*E38*^

### Adult Infections

To prepare infection vials, bacterial pellets were dissolved in filter sterilized 5% sucrose/PBS. Chromatography paper (Fisher, Pittsburgh, PA) discs were dipped in the bacterial solution (5% sucrose was used as a control) and were carefully placed on standard fly food vials such that they covered the entire food surface. Adult females were first subjected to a 2-hr starvation in empty vials at 29°C. Then 10-12 flies were transferred to each infection vial and then placed in a 29°C incubator for the duration of the assay.

### Adult survival assay

Adult female flies were infected as above. Post infection the flies were transferred to fresh food vials every 2 days. The number of deaths was scored every 24hrs.

### Rapamycin and chemical feeding

For treatment with rapamycin, 3-5 day old female flies were shifted on vials containing 200μM rapamycin dissolved in standard Drosophila food for 24hrs. DMSO dissolved in food was used as a control. After 24hr of rapamycin pre-treatment, the flies were then transferred to infection vials mixed with 200μM final concentration of rapamycin or DMSO. Chemical intestine stressors, [25μg/ml Bleomycin (Sigma # 9041-93-4), 5%DSS (Sigma, #9011-18-1) 2mM paraquat (Sigma, #75365-73-0)] were used to induce intestine specific stress in *w*^*1118*^ flies. 5% sucrose solution was used as a solvent for all the mentioned chemical stressors. 5% Sucrose solution alone was used as the control for all experiments. 500μl of each solution was used to completely soak a piece of 2.5 cm × 3.75 cm chromatography paper (Fisher, Pittsburgh, PA), which was then placed inside an empty vial). 5–7-day old, mated females (n=10-12/ vial) were then added to the vials.

### SDS-PAGE and western Blotting

Intestines (10 per sample) were dissected in ice cold 1X PBS and immediately lysed in ice cold lysis buffer containing 20 mM Tris-HCl (pH 8.0), 137 mM NaCl, 1 mM EDTA, 25 % glycerol, 1% NP-40, 50 mM NaF, 1 mM PMSF, 1 mM DTT, 5 mM sodium ortho vanadate (Na_3_VO_4_) and Protease Inhibitor cocktail (Roche Cat. No. 04693124001) and Phosphatase inhibitor (Roche Cat. No. 04906845001). Protein concentrations were measured using the Bio-Rad Dc Protein Assay kit II (5000112). Protein lysates (15 μg to 30μg) were resolved by SDS–PAGE and transferred to a nitrocellulose membrane, and then subjected to western blotting with specific primary antibodies and HRP-conjugated secondary antibodies, and then visualized by chemiluminescence (enhanced ECL solution (Perkin Elmer). Primary antibodies used in this study were: anti-phospho-S6K-Thr398 (1:1000, Cell Signalling Technology #9209), anti-pERK T980 (Cell signalling technology #3179, 1:1000 dilution), anti-pAkt-S505 (Cell Signalling #4054, 1:1000 dilution), anti-phospho S6 (gift from Aurelio Teleman) and anti-actin (1:1000, Santa Cruz Biotechnology, # sc-8432). Secondary antibodies were purchased from Santa Cruz Biotechnology (sc-2030, 2005, 2020, 1:10,000 dilution).

### Immunostaining

The fly intestines were dissected in ice cold 1X PBS. The samples were then fixed in 4% Paraformaldehyde in 1X PBS (1:4 diluted from Pierce™ 16% Formaldehyde Cat # 28906) at room temperature for 30 mins. Post fixation, the tissues were washed with 1X PBS + 1% TritonX100, for 10 mins. The tissues were then blocked in 1X PAT buffer + 2% fetal bovine serum (FBS) for 2hrs. The tissues were then transferred to fresh PAT containing the primary antibody, overnight at 4°C. The primary antibody incubation was followed by 3 washes with 1X PBT + 2% fetal bovine serum (FBS) for 30 mins each. The tissues were then incubated with secondary antibody in PBT without the serum followed by washing thrice with PBT without serum. Finally, the tissues were incubated with 1:10000dil of Hoechst 33342 (Invitrogen) in PBT to stain the nuclei. The tissues were then mounted on glass slides with coverslips, using Vectashield (Vector laboratories Inc., CA). The slides were visualized under a Zeiss Observer Z1 microscope using the 10x and 20x objectives and with Zen-Axiovision software.

### Quantitative Real Time Polymerase Chain Reaction (qRT-PCR)

Total RNA was extracted from groups of 5 adults or 10 intestines using TRIzol reagent according to manufacturer’s instructions (Invitrogen; 15596–018). The RNA samples were treated with DNase (Ambion; 2238 G) and then reverse transcribed using Superscript II (Invitrogen; 100004925). The cDNAs were then used as a template for subsequent qRT–PCRs using SyBr Green PCR mix and an ABI 7500 real time PCR system. The PCR data were normalized to actin mRNA levels.

### Bodipy staining

The adult intestines were dissected in ice cold 1X PBS. The tissue samples were then fixed in 4% Paraformaldehyde in 1X PBS (1:4 diluted from Pierce™ 16% Formaldehyde Cat # 28906) at room temperature for 30 mins. The fixation was followed by a couple washes with ice cold PBS. The BODIPY (Invitrogen) was diluted in PBS (1:100) for 30mins at RT. The samples were then washed twice with PBS for 10min. Finally, the tissues were incubated with 1:10000dil of Hoechst 33342 (Invitrogen) in PBT to stain the nuclei, followed by another wash with PBS for 10mins at RT. The tissues were then mounted on slides and visualized as mentioned above.

### Oil Red O staining

The adult intestines were dissected in ice cold PBS. The tissue samples were then fixed in 4% paraformaldehyde in PBS at RT for 30 mins. The fixation was followed by a couple washes with PBS. The intestines were then incubated in fresh Oil Red O (Sigma-Aldrich Cat # O0625) solution. The solution was prepared by adding 6ml of 0.1% Oil Red O in Isopropanol and 4ml ultra-pure dH_2_O, passed through 0.45μm syringe), followed by rinsing with distilled water. The tissues were mounted on a glass slide and the tissues were imaged using a dissecting microscope.

### TAG and glycogen assays

The metabolic assays were performed as previously described (Tennessen et al., 2014). Briefly, animals were lysed, and lysates were heated at 70 Celsius for 10 minutes. Then they were incubated first with triglyceride reagent (Sigma; T2449) and then mixed with free glycerol reagent (Sigma; F6428). Colorimetric measurements were then made using absorbance at 540 nm and TAG levels calculated by comparing with a glycerol standard curve. Glycogen assays were performed by lysing animals in PBS and then heating lysates at 70 Celsius for 10 minutes. For each experimental sample, duplicate samples were either treated with amyloglucosidase (Sigma A1602) to breakdown glycogen intro glucose, or left untreated, and then levels of glucose in both duplicates measured by colorimetric assay following the addition of a glucose oxidase reagent (Sigma; GAGO-20). Levels of glycogen in each experimental sample were then calculated by subtracting the glucose measurements of the untreated duplicate from the amyloglucosidase-treated sample. All experimental metabolite concentrations were calculated by comparison with glycogen and glucose standard curves.

## ACKNOWLEDGEMENTS

We thank Edan Foley for the gift of fly stocks. Stocks obtained from the Bloomington Drosophila Stock Center (NIH P40OD018537) were used in this study. This work was supported by a CIHR project grant and NSERC discovery grant to S.S.G..

---

**Suppl Figure 1.**
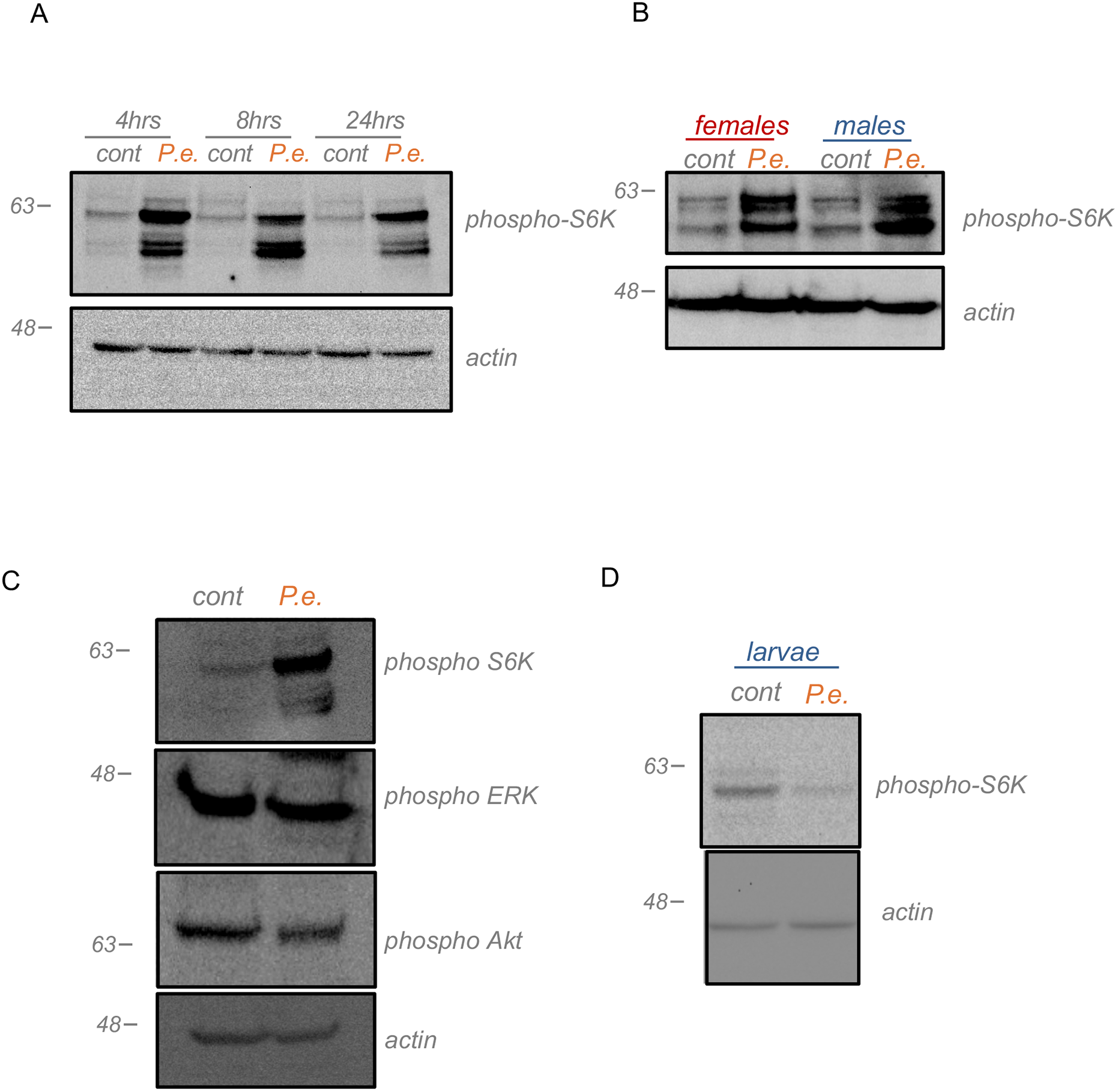
Enteric bacterial infection stimulates TOR activity in adult intestines. A) Time course of *P*.*e*. infection on phosphorylated-S6K levels in intestines of *w*^*1118*^ mated females. Dissected intestines were collected at 4hrs, 8hrs, and 24hrs post infection, lysed, and analysed by western blotting using antibodies against phosphorylated*-*S6K and actin (loading control). B) Adult *w*^*1118*^ mated flies sorted by sex and subjected to oral *P*.*e*. infection. After 4hrs, dissected intestines were lysed and processed for western blotting using antibodies against phosphorylated-S6K *and actin* (loading control). C) Adult *w*^*1118*^ mated females were subjected to 4hr oral *P*.*e*. infection. Intestinal samples were then processed for western blotting using antibodies against-phosphorylated-S6K, phosphorylated-ERK, phosphorylated-Akt, and actin. D) Third instar larvae (96 hr AEL) subjected to 4hr *P*.*e*. infection. Dissected larval intestines were lysed and processed for western blotting using antibodies against phosphorylated-S6K and actin. A representative blot is shown.

